# Citation needed? Wikipedia bibliometrics during the first wave of the COVID-19 pandemic

**DOI:** 10.1101/2021.03.01.433379

**Authors:** Omer Benjakob, Rona Aviram, Jonathan Sobel

**Affiliations:** The Cohn Institute for the History and Philosophy of Science and Ideas, Tel Aviv University, Tel Aviv, Israel; Weizmann Institute of Science, Rehovot, Israel; Faculty of Biomedical Engineering, Technion-IIT, Haifa, Israel

**Keywords:** COVID-19, Wikipedia, Infodemic, sources, bibliometrics, citizen science, open science

## Abstract

**Background:** With the COVID-19 pandemic’s outbreak, millions flocked to Wikipedia for updated information. Amid growing concerns regarding an “infodemic”, ensuring the quality of information is a crucial vector of public health. Investigating if and how Wikipedia remained up to date and in line with science is key to formulating strategies to counter misinformation. Using citation analyses, we asked: which sources informed Wikipedia’s COVID-19-related articles before and during the pandemic’s first wave (January-May 2020).

**Results:** We found that coronavirus-related articles referenced trusted media sources and high-quality academic research. Moreover, despite a surge in COVID-19 preprints, Wikipedia had a clear preference for open-access studies published in respected journals and made little use of preprints. Building a timeline of English COVID-19 articles from 2001-2020 revealed a nuanced trade-off between quality and timeliness. It further showed how preexisting articles on key topics related to the virus created a framework for integrating new knowledge. Supported by a rigid sourcing policy, this “scientific infrastructure” facilitated contextualization and regulated the influx of new information. Lastly, we constructed a network of DOI-Wikipedia articles, which showed the shifting landscape of pandemic-related knowledge on Wikipedia and how academic citations create a web of shared knowledge supporting topics like COVID-19 vaccine development.

**Conclusions:** Understanding how scientific research interacts with the digital knowledge-sphere during the pandemic provides insight into how Wikipedia can facilitate access to science. It also reveals how, aided by what we term its “citizen encyclopedists”, it successfully fended off COVID-19 disinformation and how this unique model may be deployed in other contexts.

## Introduction

Wikipedia has over 130,000 different articles relating to health and medicine (1). The website as a whole, and specifically its medical and health articles, like those about disease or drugs, are a prominent source of information for the general public (2). Studies of readership and editorship of health articles reveal that medical professionals are active consumers of Wikipedia and make up roughly half of those involved in editing these articles in English (3, 4). Research conducted into the quality and scope of medical content deemed Wikipedia “a key tool for global public health promotion” (4, 5) and others have found that in terms of content errors Wikipedia is on par with academic and professional sources even in fields like medicine (6). Meanwhile, a metastudy of all the research about Wikipedia’s health and medical content found it to be a prominent health information resource for experts and non-experts alike. (7). With the WHO labeling the COVID-19 pandemic an “infodemic” (8), and disinformation threatening public health, a closer examination of Wikipedia and its references during the pandemic is merited. Wikipedia’s “COVID-19 pandemic” article was among the most viewed in 2020 (9) - with a peak interest during the first wave. Researchers from different disciplines have looked into citations in Wikipedia and done bibliometric analyses of it - for example, asking if open-access papers are more likely to be cited in Wikipedia (10). While anecdotal research has shown that Wikipedia and its academic references can mirror the growth of a scientific field (11), few have researched the coronavirus and Wikipedia. This research has shown both that traffic to Wikipedia’s coronavirus articles reflected public interest in the pandemic (12), and that these articles provide a representative sample of COVID-19 research (13). However, to our knowledge, no research has yet focused on the pandemic’s “bibliometrics” on Wikipedia, and addressed the different dynamics regarding its sources - be they popular or academic - before and during the pandemic’s first wave.

The aim of the present study is to provide a comprehensive bibliometric analyses of english Wikipedia’s COVID-19 articles during this period. To characterize the scientific literature as well as general media sources supporting the encyclopedia’s coverage of the COVID-19 as the pandemic spread we performed citation analyses of the references used in Wikipedia’s coronavirus articles. We did this along three axes: the references used in the relevant articles at the end of the first wave, their historical trajectory, and their network interaction with Wikipedia articles on this topic.

## Material and Methods

Using citations as a metric for gauging the scientificness of Wikipedia articles along these three axes allowed us to provide a “scientific score” (1) for them and ask: what shifts in scientificness did the COVID-19 articles undergo during the period researched. At the level of the citations inside any given Wikipedia article, we could provide a second metric, namely the latency (2) which allowed us to get a historical perspective on the scientific infrastructure supporting them. Moreover our work explored Wikipedia’s articles’ versions history and co-citations, to gain an insight on COVID-19 knowledge and its growth since the creation of the digital encyclopedia in 2001 and up until 2020. Though predominantly quantitative, for some selected articles we also examined the different claims the citations were used to support, at different stages, and reviewed some of the textual changes that articles underwent in wake of the coronavirus outbreak, to provide anecdotal qualitative context to our findings.

### Corpus Delimitation

Digital Object Identifiers (DOIs) were used to identify academic sources among the references found within any given Wikipedia article. To delimit the corpus of Wikipedia COVID-19-articles containing DOIs, two different strategies were applied (Supplementary figure S1A). Every Wikipedia article affiliated with the official WikiProject COVID-19 task force (more than 1,500 pages during the period analyzed) was scraped using an R package specifically developed for this study, *WikiCitationHistoRy*. In combination with the *WikipediR* R package, which was used to retrieve the list of actual articles covered by the COVID-19 project, our *WikiCitationHistoRy* R package was used to extract DOIs from their text and thereby identified Wikipedia pages containing academic citations, termed “Wikipedia articles” in the present study. While “articles” is used for Wikipedia entries, “papers” is used to denote academic studies referenced on Wikipedia articles. Simultaneously, we also searched the EuroPMC database, using *COVID-19*, using *SARS-CoV2*, *SARS-nCoV19* as keywords to detect scientific studies published about this topic. Thus, 30,000 peer-reviewed papers, reviews, and preprint studies were retrieved. This set was compared to the DOI citations extracted from the entirety of the English Wikipedia dump of May 2020 (~860,000 DOIs) using *mwcite*. Thus, Wikipedia articles containing at least one DOI citation were identified - either from the EuroPMC search or through the specified Wikipedia project. The resulting “COVID-19 corpus” comprised a total of 231 Wikipedia articles - all related to COVID-19 and based on at least one academic source. In this study, the term “corpus” describes this body of Wikipedia “articles”, and “sets” is used to describe “papers” and the bibliographic information relating to academic studies (i.e. DOIs).

### DOI Corpus Content Analysis and DOI Sets Comparison

The analysis of DOIs led to the categorization of three DOI sets: 1) the COVID-19 Wikipedia set, 2) the EuroPMC 30K search and 3) the Wikipedia dump of May 2020. For the dump and the COVID sets, the latency was computed (to gauge how much time had passed from an academic paper’s publication until it was cited on Wikipedia), and for all three sets we retrieved their scientific citations count (the number of times the paper was cited in scientific literature), their Altmetric score, as well as the papers’ authors, publishers, journal, source type (preprint server or peer-reviewed publication), open-access status (if relevant), title and keywords. In addition, in the COVID-19 Wikipedia corpus the DOI set’s citation count among wikipedia articles were also analysed to help gauge the importance of the sources.

### Text Mining, Identifier Extraction and Annotation

From the COVID-19 corpus, DOIs, PMIDs, ISBNs, and URLs (Supplementary figure S1B) were extracted using a set of regular expressions from our R package. Moreover *WikiCitationHistoRy* allows the extraction of other sources such as tweets, press releases, reports, hyperlinks and the *protected* status of Wikipedia pages (on Wikipedia, pages can be locked to public editing through a system of “protected” statuses). Subsequently, several statistics were computed for each Wikipedia article and information for each of their DOI were retrieved using *Altmetrics* (14), *CrossRef* (15) and the *EuroPMC* (16) R packages.

### Visualisations and Metrics

Our R package was developed in order to retrieve any Wikipedia article and its content, both in the present - i.e article text, size, reference count and users - and in the past - i.e. timestamps, revision IDs and the text of earlier versions. This package allows the retrieval of the relevant information in structured tables and helped support several visualisations for the data. Notably, two navigable visualisations were created and are available for any set of Wikipedia articles: 1) A timeline of article creation dates which allows users to navigate through the growth of Wikipedia articles related to a certain topic over time, and 2) a network linking Wikipedia articles based on their shared academic references. The package also includes a proposed metric to assess the scientificness of a Wikipedia article. This metric, called *Sci Score* (shorthand for scientific score), is defined by the ratio of academic as opposed to non-academic references any Wikipedia article includes, as such:

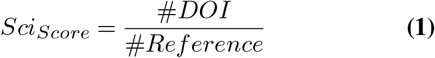

Our investigation also included an analysis of the latency (11) of any given DOI citation on Wikipedia. This metric is defined as the duration (in years) between the date of publication of a scientific paper and the date of introduction of the DOI into a specific Wikipedia article, as defined below:

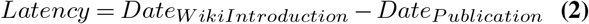

### Data and Code Availability Statement

Every table and all of our raw data are available online through the ZENODO repository with DOI: 10.5281/zenodo.3901741. Every visualisation and statistics were completed using R statistical programming language (R version 3.5.0). A beta version of the visualizations, their code and the documentation from our R package are available on the Github repositories: https://github.com/jsobel1/WikiCitationHistoRy https://github.com/jsobel1/Wiki_COVID-19_interactive_network https://github.com/jsobel1/Interactive_timeline_wiki_COVID-19

## Results

### COVID-19 Wikipedia Articles: Well-Sourced but Highly Selective

We set out to characterize the representation of COVID-19-related research on Wikipedia. As all factual claims on Wikipedia must be supported by “verifiable sources” (17), we focused on articles’ references to ask: What sources were used and what was the role of scientific papers in supporting coronavirus articles on Wikipedia? For this aim, we first identified the relevant Wikipedia articles related to COVID-19 (Supplementary figure S1A) as described in detail in the methods section. Then, we extracted relevant information such as identifiers (DOI, ISBN, PMID), references and hyperlinks (Supplementary figure S1B).

From the perspective of Wikipedia, though there were over 1.5K (1,695) COVID-19-related articles, only 149 had academic sources. We further identified an additional 82 Wikipedia articles that were not part of Wikipedia’s organic set of coronavirus articles, but had at least one DOI reference from the EuroPMC database - which consisted of over 30,000 COVID-19 related papers (30,720) (Supplementary figureS1C). Together these 231 Wikipedia articles served as the main focus of our work as they form the scientific core of Wikipedia’s COVID-19 coverage. This DOI-filtered COVID-19 corpus included articles on scientific concepts, genes, drugs and even notable people who fell ill with coronavirus. The articles ranged from “Severe acute respiratory syndrome-related coronavirus”, “Coronavirus packaging signal” and “Acute respiratory distress syndrome”, to “Charles, Prince of Wales”, “COVID-19 pandemic in North America,” and concepts with social interest like “Herd immunity”, “Social distancing”, “Wet market” or even public figures like “Dr. Anthony Fauci”. This corpus included articles that were purely about scientific topics as well as those that had both scientific and social content and that were on topics with general interest to the public. For example, the article for “Coronavirus”, the drugs “Chloroquine” and “Favipiravir,” and other less scientific articles with wider social interest, like the article for “Social distancing” and “Shi Zhengli”, the virologist employed by the Wuhan Institute of Virology and who earned public notoriety for her research into the origins of COVID-19.

Comparing the overall corpus of academic papers dealing with COVID-19 to those cited on Wikipedia we found that less than half a percent (0.42%) of all the academic papers related to coronavirus made it into Wikipedia (Supplementary figure S1C). Thus, our data reveals Wikipedia was highly selective in regards to the existing scientific output dealing with COVID-19 (See supplementary dataset (1)).

We next analyzed all the citations and references included in the complete Wikipedia dump from May 2020, using *mwcite*. Thus, we could extract a total number of about 2.68 million citations (2,686,881) comprising ISBNs, DOIs, arXiv, PMID and PMC numbers (Supplementary figure S1D). Among the citations extracted were 860K DOIs and about 38K preprints IDs from arXiv, about 1.4 percent of all the citations in the dump, indicating that the server hosting non-reviewed studies does contribute sources to Wikipedia alongside established peer-review journals. These DOIs were used as a separate group that was compared with the EuroPMC 30K DOIs (30,720) and the extracted DOIs (2,626 unique DOIs) from our initial Wikipedia COVID-19 set in a subsequent analysis.

An analysis of the journals and academic content from the 2,626 DOIs that were cited in the Wikipedia COVID-19 corpus reveals a strong bias towards high impact factor journals in both science and medicine. For example, *Nature* - which has an impact factor of over 42 - was among the top cited journals, alongside *Science*, *The Lancet* and the *New England Journal of Medicine*; together these four comprised 13 percent of the overall academic references (Figure 1A). The Cochrane database of systematic reviews was also among the most cited academic sources (WPM and Cochrane have an official partnership). Notably, the papers cited tended tonot just to come from high impact factor journals, but also have a higher Altmetric score compared to the overall average of papers cited in Wikipedia in general. In other words, the papers cited on Wikipedia’s COVID-19 articles were not just academically respected, but were also popular - i.e. they were shared extensively on social media such as Twitter and Facebook.

**Fig. 1.**
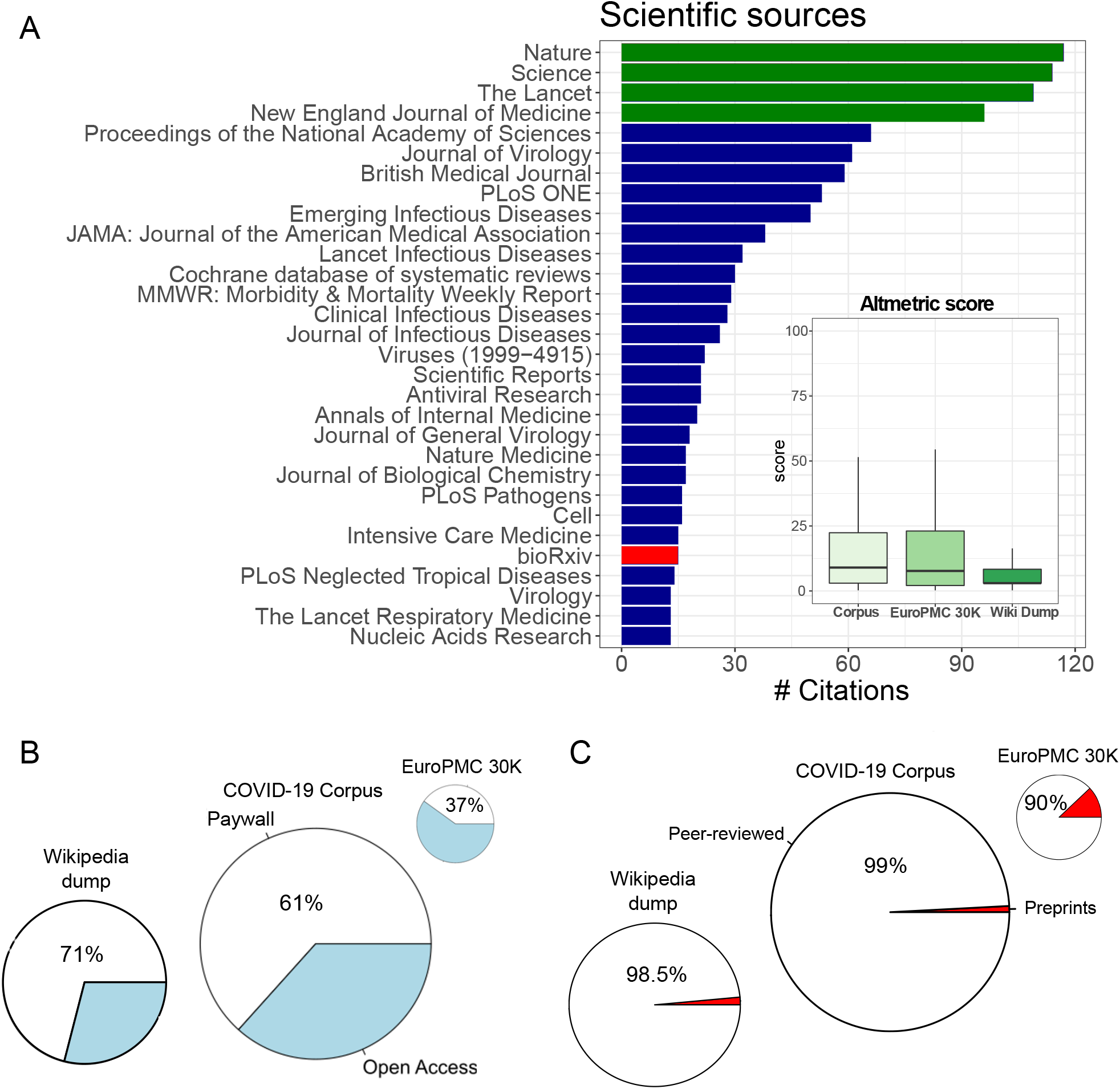
Wikipedia COVID-19 Corpus of scientific sources reveals a greater fraction of open-access papers as well as a higher impact in Altmetric score. A) Bar plot of the most trusted academic sources. Top journals are highlighted in green and preprints are represented in red. Bottom right: boxplot of the distribution Altmetrics score in Wikipedia COVID-19 corpus - the dump from May 2020, the COVID-19 Corpus and the scientific sources from the Europmc COVID-19 search. B) Fraction of open-access sources, C) fraction of preprints from bioRxiv and medRxiv.

Most importantly perhaps, we also found that more than a third of the academic sources (39%) referenced in COVID-19 articles on Wikipedia were open-access papers (Figure 1B). The relation between open-access and paywalled academic sources is especially interesting when compared to Wikipedia’s references writ large: About 29 percent of all academic sources on Wikipedia are open-access, compared to 63 percent in the COVID-19-related scientific literature (i.e. in EuroPMC).

Remarkably, despite a surge in COVID-19 research being uploaded to preprint servers, we found that only a fraction of this new output was cited on Wikipedia - less than 1 percent, or 27 (Figure 1C, Table S1) bioRxiv or medRxiv preprints were referenced. Among the preprints that were cited on Wikipedia was an early study on *Remdesivir* (18), a study on the mortality rate of elderly individuals (19), research on COVID-19 transmission in Spain (20) and New York (21), and research into how Wuhan’s health system managed to eventually contain the virus (22), showing how non-peer-reviewed studies touched on medical, health and social aspects of the virus. The later was especially prevalent with two of the preprints focusing on the benefits of contact tracing (23, 24). The number of overall preprints was in line with the general representation of preprints in Wikipedia (1.5%), but lower than would be expected considering the fact that our academic database of EuroPMC papers had almost 3,700 preprints - 12.3 percent of the roughly 30,000 COVID-19 related papers in May 2020. Thus, in contrast to the high enrichment of preprints in COVID-19 research, Wikipedia’s editors overwhelmingly preferred peer-reviewed papers to preprints. In other words, Wikipedia generally cites preprints more than it was found to on the topic of COVID-19, while COVID-19 articles cited open-access paper by more 10% (from 29% to 39%). Taken together with the bias towards high-impact journals, our data suggest that this contributed significantly to Wikipedia’s ability to stay both up to date and to maintain high academic standards, allowing editors to cite peer-reviewed research despite other alternatives being available.

Due to the high selectivity of Wikipedia editors in terms of the percentage of COVID-19 academic research actually cited on Wikipedia’s COVID-19 articles, we also focused on non-academic sources. Popular media, we found, played a substantial role in our corpus. Over 80 percent of all the references used in the COVID-19 corpus were non-academic, being either general media or websites (Figure 2A). In fact, a mere 16 percent of the over 21,000 references supporting the COVID-19 content were from academic journals. Among the general media sources used (Figure 2B–D), there was a high representation for what is termed legacy media outlets, like the *New York Times* and the *BBC*, alongside widely syndicated news agencies like *Reuters* and the *Associated Press*, and official sources like *WHO.org* and *gov.UK*. Among the most cited websites, for example, there was an interesting representation of local media outlets from countries hit early and hard by the virus, with the Italian *La Republica* and the Chinese *South China Post* being among the most cited sites. The World Health Organization was one of the most cited publisher in the corpus of relevant articles, more than 150 references.

**Fig. 2.**
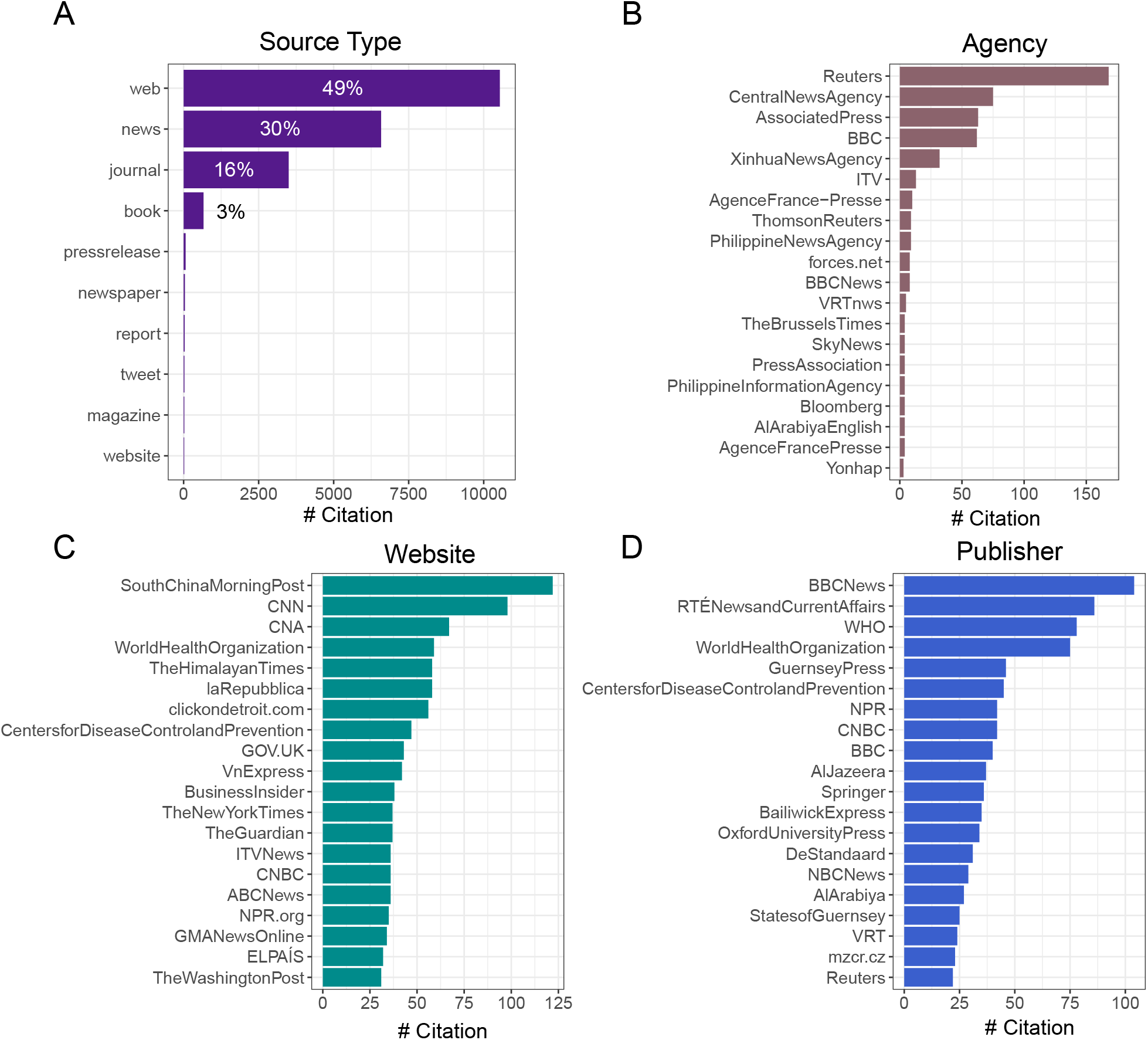
Wikipedia COVID-19 Corpus: Non-scientific sources mostly referred to websites or news media outlets considered highly respected and deemed to be trusted sources, including official sources like the WHO. A) source types extracted from the COVID-19 corpus of Wikipedia articles B) most cited news agencies, C) most cited websites and D) most cited publisher form the COVID-19 Corpus sources

### Scientific Score

To distinguish between the role scientific research and popular media played, we created a “scientific score” for Wikipedia articles (1). The metric is based on the ratio of academic as opposed to non-academic references any article includes. This score attempts to rank the *scientificness* of any given Wikipedia article based solely on its list of references. Ranging from 1 to 0, an article’s scientific score is calculated according to the ratio of its sources that are academic (i.e. contain DOIs), so that an article with a score of 1 will have 100 percent academic references, while that with none will have a score of zero. Technically, as all of our corpus of coronavirus-related Wikipedia articles had at least one academic source in the form of a DOI, their scientific scores will always be greater than zero (Supplementary Figure S2, Supplementary Figure S5C).

In effect, this score puts forth a metric for gauging the prominence of academic texts in any given article’s reference list - or lack thereof. Out of our 231 Wikipedia articles, 15 received a perfect scientific score of 1 (Supplementary Figure S2A). High scientific score Wikipedia articles included the articles for the enzymes of “Furin” and “TMPRSS2” - whose inhibitor has been proposed as a possible treatment for COVID-19; “C30 Endopeptidase” - a group of enzymes also known as the “SARS coronavirus main proteinase”; and “SHC014-CoV” - a form of COVID-19 that affects the Chinese rufous horseshoe bat.

In contrast to the articles with scientific topics and even biographical articles about scientists themselves, which both had high scientific scores, those with the lowest scores (Supplementary Figure S2B) seemed to focus almost exclusively on social aspects of the pandemic and its immediate outcome. For example, the articles with the lowest scores dealt directly with the pandemic in a hyper-local context, including articles about the pandemic in Canada, North America, Indonesia, Japan or even Jersey, to name a few. Others focused on different aspects of the pandemic, for example the “Impact of the COVID-19 pandemic on the arts and cultural heritage” or “Travel restrictions related to the COVID-19 pandemic”. One of the articles with the lowest scientific score was the “Trump administration communication during the COVID-19 pandemic” which made scarce use of coronavirus-related research to inform its content, citing a single academic paper (related to laws regulating quarantine) among its 244 footnotes.

### The Price of Remaining Up to Date on COVID-19

During the pandemic, there were over tens of thousands of edits to the site, with thousands of new articles being created and scores of existing ones being re-edited and recast in wake of new developments. Therefore, one could expect a rapid growth of articles on the topic, as well as a possible overall increase in the number of citations of all kinds. We sought to explore the temporal axis of Wikipedia’s coverage of the pandemic to see how coverage of COVID-19 developed, namely, what were the dynamics of the growth of COVID-19 articles and their academic references.

First, we laid out our corpus of 231 articles across a timeline according to each article’s respective date of creation (Supplementary Figure S3). An article count starting from 2001, when Wikipedia was first launched, and up until May 2020, shows that for many years there was a relatively steady growth in the number of articles that would become part of our corpus - until the pandemic hit, causing a massive peak at the start of 2020 (Figure 3A). As the pandemic spread, the total number of Wikipedia articles dealing with COVID-19 and supported by scientific literature almost doubled - with a comparable number of articles being created after 2020 than the entire time before (Figure 3A, Supplementary FigureS3) (from 134 before 2020 compared to 97 in 2020).

**Fig. 3.**
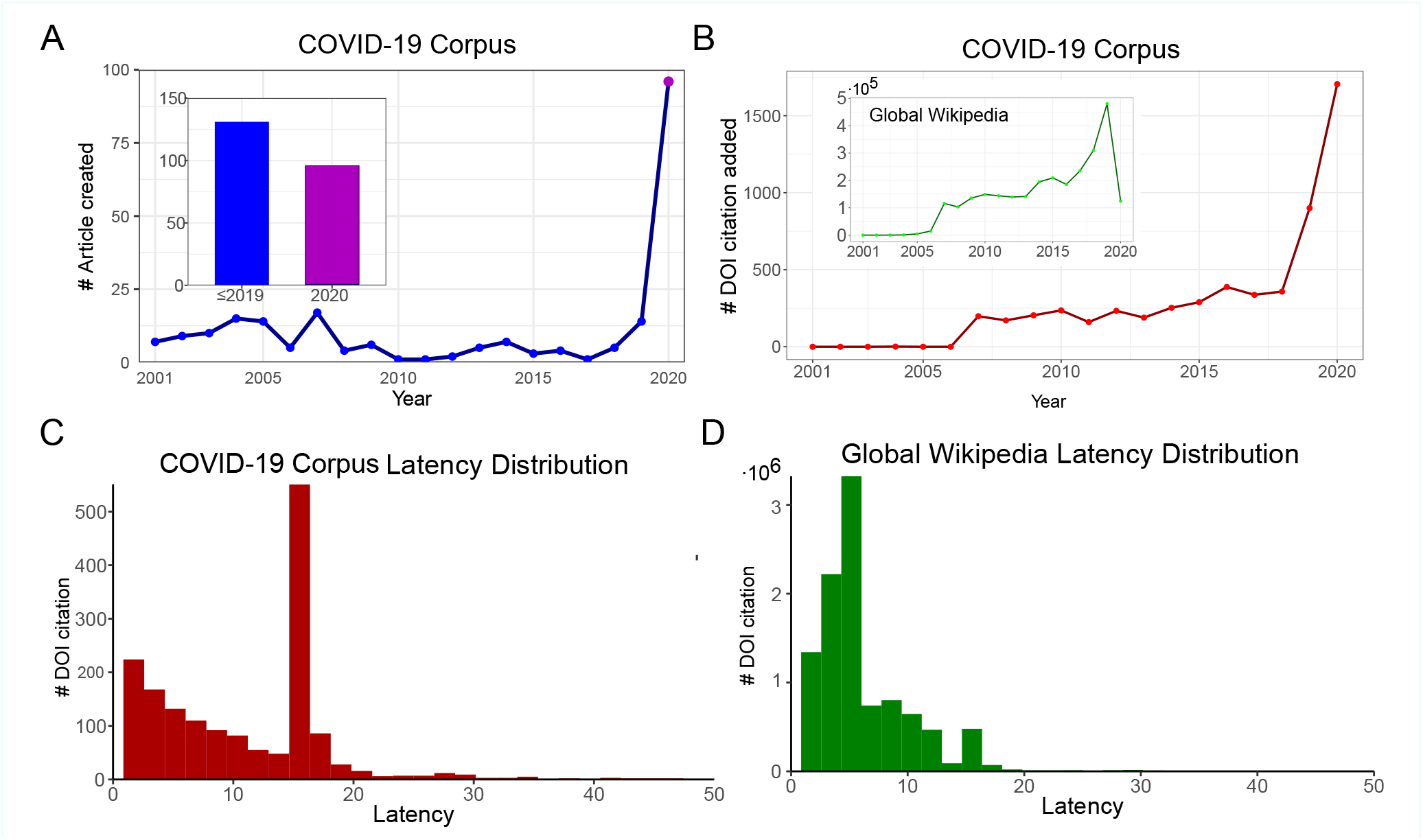
Historical perspective of the Wikipedia COVID-19 corpus outlining the growth of COVID-19 on the encyclopedia. A) COVID-19 article creation per year and number of articles created before the pandemic compared to the first five months of the pandemic. B) Scientific citation added per year in the COVID-19 category and globally in Wikipedia. C) Latency distribution of scientific literature in the COVID-19 corpus and D) latency distribution of scientific literature in the Wikipedia dump. See Supplementary FigureS3 and here for an interactive version of the timeline.

The majority of the pre-2020 articles were created relatively early - between 2003 and 2006, likely linked to a general uptick in creation of articles on Wikipedia during this period. For example, the article for (the non-novel) “coronavirus” has existed since 2003, the article for the medical term “Transmission” and that of “Mathematical modeling of infectious diseases” from 2004, and the article for the “Coronaviridae” classification from 2005. Articles opened in this early period tended to focus on scientific concepts - for example those noted above or others like “Herd immunity”. Conversely, the articles created post-pandemic during 2020 tended to be hyper-local or hyper-focused on the virus’ effects. Therefore, we collectively term the first group Wikipedia’s “scientific infrastructure”, as they allowed new scientific information to be added into existing articles, alongside the creation of new ones focusing on the pandemic’s actual ramifications.

Examining the date of publication of the peer-reviewed studies referenced on Wikipedia shows that new COVID-19 research was cited alongside papers from previous years and even the previous century, the oldest being a 1923 paper titled the “The Spread of Bacterial Infection. The Problem of Herd-Immunity.” (25). Overall, among the papers referenced on Wikipedia were highly cited studies, some with thousands of citations (Table 3), but most had relatively low citation counts (median of citation count for a paper in the corpus was 5). Comparing between a paper’s date of publication and its citation count reveals there is low anti-correlation (−0.2) but highly significant between the two (Pearson’s product-moment correlation test p-value < 10^−15^, Figure S5A). This suggest that on average older scientific papers have a higher citation count; unsurprisingly, the more time that has passed since publication, the bigger the chances a paper will be cited.

The pre-pandemic articles tended to have a high scientific score - for example, “Chloroquine”, which has been examined as a possible treatment for COVID-19 - but also underwent a shift in content in wake of the pandemic, seeing both a surge in traffic and a surge in editorial activity (Supplementary Figure S4). However, per a subjective reading of this article’s content and the editing it underwent during this period, much of the scientific content that was present pre-pandemic remained intact, with new coronavirus-related information being integrated into the existing content. The same occurred with many social concepts retroactively affiliated with COVID-19. Among these we can note the articles for “Herd immunity”, “Social distancing” and the “SARS conspiracy theory” that also existed prior to the outbreak and served as part of Wikipedia’s scientific infrastructure, allowing new information to be contextualized.

In addition to the dramatic rise in article creation during the pandemic, there was also a rise in the overall number of references affiliated with COVID-19 articles on Wikipedia (Figure 3B). In fact, the number of added DOIs in our articles grew almost six-fold post-2020 - from roughly 250 to almost 1,500 citations. Though most of the citations added were not just academic ones, with URLs overshadowing DOIs as the leading type of citation added, the general rise in citations can be seen as indicative of scientific literature’s prominent role in COVID-19 when taking into account that general trend in Wikipedia: The growth rate of references on COVID-19 articles was generally static until the outbreak; but on Wikipedia writ large references were on a rise since 2006. The post-2020 surge in citations was both academic and non-academic (Supplementary Figure S5B).

One could hypothesize that a rapid growth in the number articles dedicated to coronavirus would translate to an overall decrease in the presence of academic sources, as Wikipedia can create newer articles faster than academic research can be published on current events. Comparing the pre- and post-2020 articles’ scientific score reveals that on average, the new articles had a mean score of 0.14, compared to the pre-2020 group’s mean of 0.48 and the overall average of 0.3 (Supplementary Figure S5C). Reading the titles of the 2020 articles to glean their topic and reviewing their respective scientific score can also point to a generalization: the more scientific an article is in topic, the more scientific its references are - even during the pandemic. This means that despite the dilution at a general level during the first month of 2020, articles with scientific topics created during this period did not pay that heavy of an academic price to stay up to date.

How is that Wikipedia managed to maintain academic sourcing on new and old articles about coronavirus as the pandemic was happening? One possible explanation is that among the academic papers added to Wikipedia in 2020 were also papers published prior to this year if not a long time before. To investigate this hypothesis we used the latency metric (2). We found the mean latency of Wikipedia’s COVID-19 content to be 10.2 years (Figure 3C), slower than Wikipedia’s overall mean of 8.7 (Figure 3D). In fact, in the coronavirus corpus we observed a peak in latency of ~17 years - with over 500 citations being added to Wikipedia 17 years after their initial academic publication - almost twice as slow as Wikipedia’s average. Interestingly, this time frame corresponds to the SARS outbreak (SARS-CoV-1) in 2002-2004, which yielded a boost of scientific literature regarding coronaviruses. This suggests that while there was a surge in editing activity during this pandemic that saw papers published in 2020 added to the COVID-19 articles, a large and even prominent role was still permitted for older literature. Viewed in this light, older papers played a similar role to prepandemic articles, giving precedence to existing knowledge in ordering the integration new knowledge on scientific topics.

Comparing the articles’ scientific score to their date of creation portrays Wikipedia’s scientific infrastructure and its dynamics during the pandemic (Supplementary Figure S5C). It reveals that despite maintaining high academic standards, citing papers published in prestigious and high impact factor journals, the need to stay up to date with COVID-19 research did come at some cost: most of the highest scoring articles were ones created pre-pandemic (mostly during 2005-2010) and newer articles had a lower scientific score (Supplementary Figure S5C).

### Networks of COVID-19 Knowledge

To further investigate Wikipedia’s scientific sources and its infrastructure, we built a network of Wikipedia articles linked together based on their shared academic (DOI) sources. We filtered the list of papers (extracted DOIs) in order to keep those which were cited in at least two different Wikipedia articles, and found 179 that fulfilled this criteria, mapped to 136 Wikipedia articles in 454 different links (Figure 4, supplementary data (2)). This allowed us to map how scientific knowledge related to COVID-19 played a role not just in specific articles created during or prior to the pandemic, but actually formed a web of knowledge that proved to be an integral part of Wikipedia’s scientific infrastructure. Similar to the timeline described earlier and as a subset of our COVID-19 corpus, Wikipedia articles belonging to this network included those dealing with people, institutions, regional outcomes of the pandemic as well as scientific concepts, for example those regarding the molecular structure of the virus or the mechanism of infection (“C30 Endopeptidase”, “Coronaviridae”, and “Airborne disease“). It also included a number of articles regarding the search for a potential drug to combat the virus or other possible interventions against it (articles on topics like social distancing, vaccine development and drugs in current clinical trials).

**Fig. 4.**
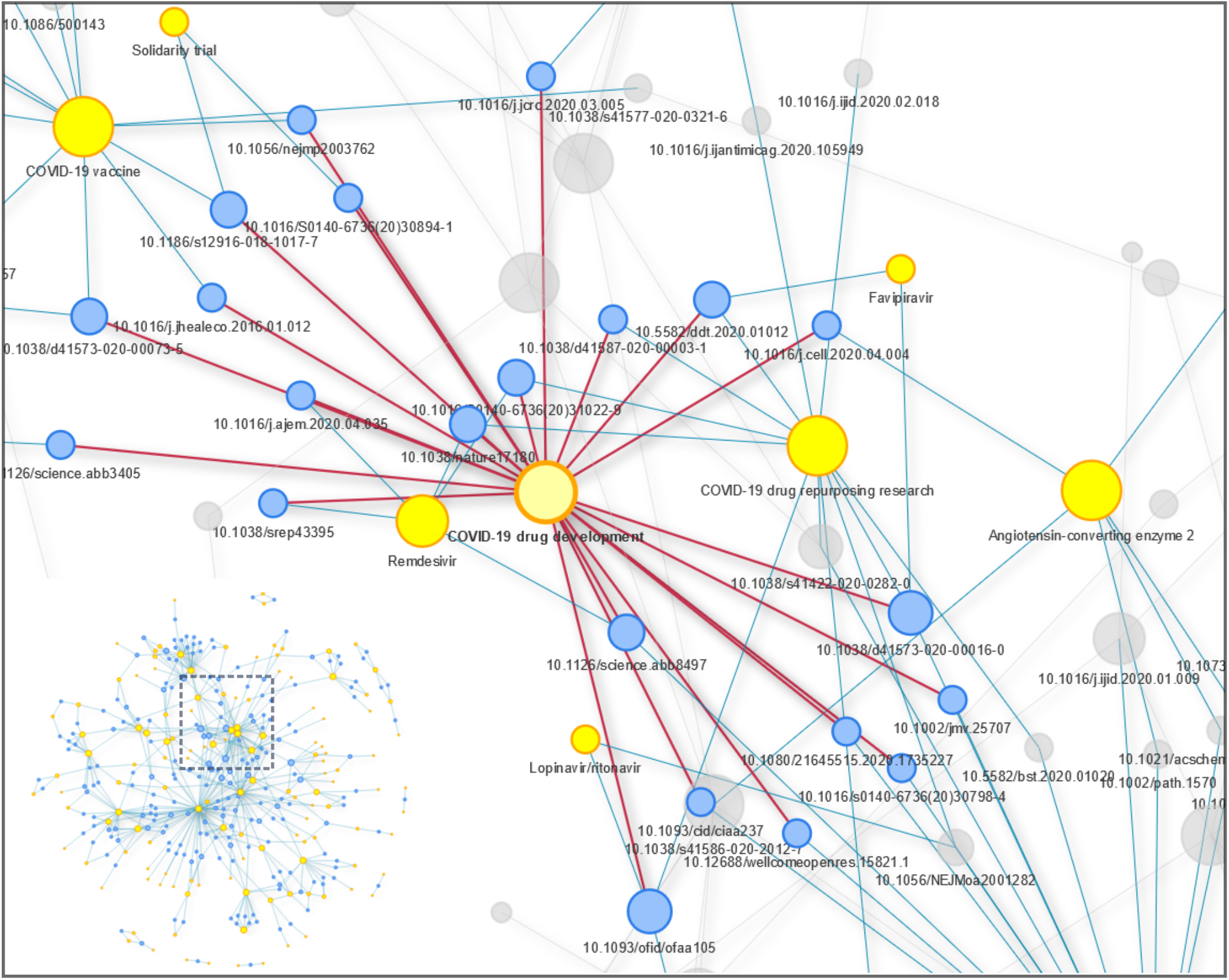
Wikipedia COVID-19 corpus article-scientific papers (DOI) network. The network mapping scientific papers cited in more than one article in the Wikipedia COVID-19 corpus was constructed using each DOI connecting at least two Wikipedia articles. This network is composed of 454 edges, 179 DOIs (Blue) and 136 Wikipedia articles (Yellow). A zoom in on the cluster of Wikipedia articles dealing with COVID-19 drug development is depicted with edges in red connecting the DOIs cited directly in the article and edges in blue connecting these DOIs to closely related articles citing the same DOIs. See here for an interactive version of the network. See Supplementary dataset (2).

Interestingly, we observed six prominent Wikipedia articles emerge in this network. These shared multiple citations with many other pages through DOI connections (nodes with an elevated degree). Four of these six so-called major nodes had a distinct and broad topic: “Coronavirus,” which focused on the virus writ large; “Coronavirus disease 2019”, which focused on the pandemic; and “COVID-19 drug repurposing research” and “COVID-19 drug development.” The first two articles were key players in how Wikipedia presented its coverage of the pandemic to readers: both were linked to from the main coronavirus article (“COVID-19 pandemic“) which was placed on the English Wikipedia’s homepage in a community-led process known as “In the News” which showcases select articles on relevant topics on the website’s homepage. Later on, alongside this community process led by the volunteers of the WikiProject COVID-19 task force, the WikiMedia Foundation also issued a directive to place a special banner referring to the “COVID-19 pandemic” article on the top of every single article in English, driving millions to the article and to subsequent articles linking out from it. These, too, were part of our network, showing how citations shared between articles can also coincide with inter-connectivity between the articles themselves.

The two remaining nodes were similar and did not prove to be distinctly independent concepts, but rather interrelated ones, with the articles for “Severe acute respiratory syndrome–related coronavirus” and “Severe acute respiratory syndrome coronavirus” each appearing as their own node despite their thematic connection. It is also interesting to note that four of the six Wikipedia articles that served as the respective centers of these groups of nodes were locked to public editing as part of the protected page status (see supplementary data (3)) and these were all articles linked to the WikiProject Medicine or, at a later stage, to the specific offshoot project set up to deal with COVID-19.

Two main themes that emerge from the network is that of COVID-19 related drugs and of the disease itself (Figure 4). Unlike popular articles relating to the effect of the virus, which we have seen are predominantly based on popular media, with scientific media playing a relatively small role, these two were topics that did require scientific basing to be able to be reliable according to the MEDRS policy - short-hand for “medical reliable sources, the sourcing policy is Wikipedia’s most rigit and bans primary sources and instead demands meta-analysis or secondary sources that provide an overview of existing research and multiple-case-study clinical trials (26). The prominence of articles like “Coronavirus disease 2019” or “COVID-19 drug development” - both of which were locked (supplementary dataset (3)) and fell under the auspices of the COVID-19 task force - in our network underscore the role academic media had in their references. Furthermore, it highlights the effects of the editing community’s centralized efforts: for example, by allowing key studies to find a role both in popular articles reached from the main articles and in scientific articles linking out from them, thus creating the network we describe.

In our network analysis, an additional smaller group of nodes (with a lower degree) was also found. It had to do almost exclusively with China-related issues. As such, it exemplified how Wikipedia’s sourcing policy - which has an explicit bias towards peer-reviewed studies and is enforced exclusively by the community - helps fight disinformation. For example, the academic paper that was most cited in Wikipedia’s COVID-19 articles was a paper published in Nature in 2020, titled “A pneumonia outbreak associated with a new coronavirus of probable bat origin” (Table S2). This paper was referenced in eight different Wikipedia articles, two among which dealt directly with scientific topics - “Angiotensin-converting enzyme 2” and “Severe acute respiratory syndrome coronavirus 2” - and two dealing with what can be termed para-scientific terms linked to COVID-19 - the “Wuhan Institute of Virology” and “Shi Zhengli”. This serves to highlight how contentious issues with a wide interest for the public - in this case, the origin of the virus - receive increased scientific support on Wikipedia, perhaps as result of editors attempting to fend off misinformation supported by lesser, non-academic sources - specifically media sources from China itself, which as we have seen were present on Wikipedia. Of the five most cited papers inside the COVID-19 corpus (Table S2) three focused specifically on either bats or the virus’ animal origins, and another focused on its spread from Wuhan, China. Interestingly, one of the 27 preprints cited (Table S1) was also the first study to suggest the virus’ origin lay with bats was (27).

Taken together with the previous findings, centralized efforts in the form of locking articles did not just allow the enforcement of a rigid sourcing policy but also created a filtered knowledge funnel of sorts, which harnessed Wikipedia’ pre-existing infrastructure of articles, mechanisms and policies to allow a regulated intake of new information as well as the creation of new articles, both based on existing research.

## Discussion

In the wake of COVID-19 pandemic, characterizing scientific research on English-language Wikipedia and understanding the role it plays is both important and timely. Millions of people - both medical professionals and the general public - read about health online (1). Research has shown traffic to Wikipedia articles follows topics covered in the news (28) - a dynamic which played out during the pandemic’s first wave (12). Moreover, scientometric research has shown that academic research follows a similar pattern - with a surge of new studies during a pandemic and then a decrease after it wanes (29). During a pandemic, as was during the Zika and SARS outbreaks (30), the risk of disinformation on Wikipedia’s content is more severe. Thus, throughout the outbreak of the COVID-19 pandemic, the threat was hypothetically increased: as a surge in traffic to Wikipedia articles, research has found, often translates into an increase in vandalism (31). Moreover, research into medical content on Wikipedia found that people who read health articles on the open encyclopedia are more likely to hover over, or even read its references to learn more about the topic (32). Particularly in the case of the coronavirus outbreak, Wikipedia’s role as such took on potentially lethal consequences as the pandemic was deemed to be an *infodemic*, and false information related to the virus was deemed a real threat to public health by the UN and WHO (8). So far, most research into Wikipedia has revolved either around the quality, readership or editorship of health content on Wikipedia - or about references and sourcing in general. Meanwhile, research on Wikipedia and COVID-19 has focused almost exclusively on editing patterns and users behaviors (12), or the representativity of academic citations (13). Therefore, we deployed a comprehensive bibliometric analyses of COVID-19-related Wikipedia articles - focusing on article’s text and source, their growth over time and their network relations.

Perhaps counter-intuitively, we found that despite the traffic surge, these articles relied on high quality sources, from both popular media and academic literature. Though the proportion of academic references in newly created articles did decrease in comparison to the period before the pandemic (resulting in lower scientific score), we found that they still played a prominent role and that high editorial standards were generally maintained, utilizing several unique solutions which we will now attempt to outline and discuss based on our findings.

One possible key to Wikipedia’s success had to do with the existence of centralized oversight mechanisms by the community of editors that could be quickly and efficiently deployed. In this case, the existence of the WikiProject Medicine - one of Wikipedia’s oldest community projects - and the formation of a specific COVID-19 task force in the form of WikiProject COVID-19, helped harness exiting editors and practices like locking articles to safeguard quality across large swaths of articles and thus enforce a relatively unified sourcing policy on articles dealing with both popular and scientific aspects of the virus.

In general, all factual claims on Wikipedia need to be supported by a verifiable source. Specifically, biomedical articles affiliated with the WikiProject Medicine (WPM) are bound by a specific policy known as MEDRS (which requires meta-analysis or secondary sources for medical content (26)). However, the mere existence of this policy does not necessarily mean it is respected. However, our findings indicate that this policy, aided by the infrastructure provided by the community to enforce it, likely played a key role in regulating the quality of coronavirus articles. One mechanism used generally by the WPM to enforce the MEDRS sourcing standards and specifically deployed by the COVID-19 task force during the pandemic was locking articles to public editing (protected pages, supplementary dataset (3)). This is a technique that is used to prevent vandalism on Wikipedia (33) and is commonly used when news events drive large amounts of new readers to specific Wikipedia articles, increasing the risk of substandard sources being added into the article by editors unversed in Wikipedia’s standards. This ad hoc measure of locking an article, deployed by a community vote on specific articles for specific amounts of time, prevents anonymous editors from being able to contribute directly to an article’s text and forces them to work through an experienced editor, thus ensuring editorial scrutiny. This measure is in line with our findings that many of the COVID-19 network central nodes were locked articles.

Another possible key to Wikipedia’s ability to maintain the WPM’s MEDRS policy of high quality sources during the pandemic was the existence of a specific infrastructure related directly to sourcing. The WPM has formed institutional-level partnerships to provide editors with access to reputable secondary sources on medical and health topics - namely through its cooperation with the Cochrane Library. The Cochran Reviews’ database is available to Wikipedia’s medical editors and it offers them access to systematic literature reviews and meta-analyses summarizing the results of multiple medical research studies(34). As well as the existence of this database on medical content, the practice of providing access to high-quality sources was also deployed specifically in regards to coronavirus in the form of the task force’s list of “trusted” sources. Alongside Cochrane studies, the WHO, for example, was given special status and preference (35). This was evident in our results as the WHO was among the most cited publishers on the COVID-19 articles. Also among the most cited scientific sources were others that were promoted by the task force as preferable sourcing for scientific content: for example, *Science*, *Nature* and *The Lancet*. This indicates that list of sources recommended by the task force were actually utilized by the volunteers and thus underscores the connection between our findings and the existence of a centralized community effort.

This was also true for non-academic sources: Among general media sources that the task force endorsed were *Reuters* and the *New York Times*, which were also prominently represented in our findings. As each new edit to any locked COVID-19 article needed to be vetted by an experienced volunteer from the task force before it could go online within the body of an article’s text, the influx of new information being added was slowed down and regulated; the source list thus allowed an especially strict sourcing policy to be rigorously implemented across thousands of articles. This was true despite the fact that there is no academic verification for volunteers - in fact, research suggests that less than half of Wikipedia’s editors focused on health and medical issues are medical professionals (3, 4) - meaning that the task forces and its list of sources allowed non-experts to enforce academic-level standards.

This dynamic was also evident within articles with purely scientific content. Despite a deluge of preprints (both in general in recent years and specifically during the pandemic (36, 37)), in our analysis, non-peer-reviewed academic sources did not play a key role on Wikipedia’s coronavirus content, while open access papers did. Therefore, one could speculate that our finding that open-access papers were disproportionately cited may provide an explanation - with academic quality trumping speed, and editors opting against preprints and preferring published studies instead. Previous research has found open-access papers are more likely to be cited on Wikipedia by 47 percent (10) and nearly one-third of the Wikipedia citations link to an open-access source (38). Here we also saw that open-access was prevalent in Wikipedia and even more so on COVID-19 articles. This, we suggest, allowed Wikipedia’s editors (expert or otherwise) to keep articles up to date without reverting tonon-peer-reviewed academic content. This, one could suggest, was likely facilitated or at least aided by the decision by academic publications’ like *Nature* and *Science* to lift paywall and open public access to all of their COVID-19-related research papers, both past and present.

In addition to the communal infrastructure’s ability to regulate the addition of new information and maintain quality standards over time, another facet we found to contribute in permitting Wikipedia to stay accurate during the pandemic is what we term its scientific i nfrastructure. R esearch on Wikipedia articles’ content has shown that the initial structuring of information on a given article tends to dictate its development in later stages, and that substantial reorganizations gradually decrease over time (39). A temporal review of our articles and their citations, showed that the best-sourced articles, those with the highest scientific score that formed the scientific backbone of Wikipedia’s COVID-19 content, were those created from 2005 and until 2010. These, we argue, are part of Wikipedia’s wider scientific infrastructure, which regulated the intake of new knowledge into Wikipedia.

Our network analysis reflects the pivotal role preexisting content played in contextualizing the science behind many popular concepts or those made popular by the pandemic. Preexisting content in the form of Wikipedia articles, policies, practices, and academic research served as a framework that helped regulate the deluge of new information, allowing newer findings to find a place within Wikipedia’s existing network of knowledge. Future work on this topic could focus on the question of whether this dynamic changed as 2020 progressed and, at a later time, on how contemporary peerreviewed COVID-19-related research that was published during the pandemic’s next waves would be integrated into these articles.

Previous research has suggested that in terms of content errors Wikipedia is on par with academic and professional sources even in fields like medicine (6). A recent metaanalysis of studies about medical content on Wikipedia found that despite the prominent role Wikipedia plays for the general public, health practitioners, patients and medical students, the academic discourse around Wikipedia within the context of health is still limited (7). This indicates that academic publications and scientists are lagging on embracing it and its benefits. Such a process could help improve Wikipedia’s content and even introduce new editors with academic background into the fold, which would further improve quality and and timeliness.

Moreover, our findings suggest that “ open” science - not just open access - may be key to understanding Wikipedia’s mechanisms and how they can be translated to other contexts. In this regard, much like citizen scientists help support institutional science (40), Wikipedia’s editors may be regarded as citizen encyclopedists (11). Viewed as such, Wikipedia’s citizen encyclopedists can play the same role communicating science that citizen scientists play in creating science. However, as previous citizen science projects have taught us (41), for that to work, citizens need scientists to provide the framework for non-expert contributions (42, 43). As this study shows, a similar infrastructure can be seen to exist on Wikipedia for encyclopedic as opposed to scientific work. Thus, should the cooperation between the scientific and Wikipedia communities increase, it could be utilized for other contexts as well.

Our findings outline ways in which Wikipedia managed to fend off disinformation and stay up to date. With Facebook and other social media giants struggling to implement both technical and human-driven solutions to disinformation from the top down, it seems Wikipedia’s dual usage of established science and a community of volunteers, provides a possible model for how this can be achieved - a valuable goal during an infodemic. Some have already suggested that the American Center for Disease Control should adopt Wikipedia’s model to help communicate medical knowledge (44). In October 2020, the WHO and Wikimedia, the foundation that oversees the Wikipedia project, announced they would cooperate to make critical public health information available. This means that in the near future, the quality of Wikipedia’s coverage of the pandemic will very likely increase just as its role as central node in the network of knowledge transference to the general public becomes increasingly clear.

Wikipedia’s main advantage is in many ways its largest disadvantage: its open format which allows a large community of editors of varying degrees of expertise to contribute. This can lead to large discrepancies in article quality and inconsistencies in the ways editors add references to articles’ text (38). We tried to address these limitations using technical solutions, such as regular expressions for extracting URLs, hyprelinks, DOIs and PMIDs. In this study, which was limited to English, we retrieved most of our scientific literature metadata using Altmetrics (14, 45), EuroPMC (16) and CrossRef (15) R APIs. However the content of the underlying databases is not always accurate, and at a technical level, this method was not without limitations. For example, we could not retrieve all of the extracted DOIs’ metadata. Moreover, information regarding open access (among others) varied with quality between the APIs (46). In addition, our preprint analysis was mainly focused on MedRxiv and BioRxiv which have the benefit of having a distinct DOI prefix. Unfortunately, we found no better solution to annotate preprints from the extracted DOIs. Preprint servers donot necessarily use the DOI system (47) (i.e ArXiv) and others share DOI prefixes with published paper (for instance the preprint server used by The Lancet). Moreover, we developed a parser for general citations (news outlets, websites, publishers), and we could not avoid redundant entries (i.e “WHO”, “World Health Organisation”).

In addition, our method to delimitate the COVID-19 corpus focused on medical content (EuroPMC search) and may explain why we found predominately biomedical and health studies. However, using DOI filtering on Wikipedia’s corona virus articles should have equally led us to find studies from the social sciences - should those have been used. However, it seems that as these socially focused articles donot fall under the MEDRS sourcing policy, there was less if any use of academic studies, resulting in a low scientific score, thus highlighting the importance of this policy in enforcing academic standards on the open encyclopedia’s articles.

Finally, as Wikipedia is constantly changing, some of our conclusions are bound to change. Therefore, our study, though limited, is focused on the pandemic’s first wave and its history on English Wikipedia alone, a crucial arena for examining the dynamics of knowledge online at this pivotal time frame. As these findings regarding the first wave were the result of a robust community effort that utilized English Wikipedia’s policies and mechanisms to safeguard existing content and regulate the creation of new content, it may be specific to English Wikipedia and its community.

However, it seems safe to speculate that at least on English Wikipedia the processes will continue to take place in the future as new textual additions are made to the open encyclopedia. In fact, one could suggest that as more time passes from the first wave, the newer post-pandemic articles that had low scientific scores w ill u ndergo a r eview and h ave their sources improved as newer research becomes more readily available. Studying the second wave - for example, shifts in the scientific score overtime - and understanding how encyclopedic content written during the first wave changed over the next year could very instructive. Analyses of coronavirus articles indicated that at least on medical and health topics - especially those in the news and driving public interest - Wikipedia’s methods for safeguarding its standards withstand the test. Perhaps as more academic research regarding the virus passes review and is published in 2021 and in the coming years, the ability of Wikipedia to reduce latency on this topic without having to compromise its scientificness will increase. Moreover, our findings hint that should journals open access to research in other fields, it may help Wikipedia cite even more peer reviewed research instead of media sources or preprints. Thus, with the help of community enforcement, like that seen during the first wave of the pandemic, Wikipedia should be able to succeed in other fields as well.

In summary, our findings reveal a trade off between timeliness and scientificness in regards to scientific literature: most of Wikipedia’s COVID-19 content was supported by references from highly trusted sources - but more from the general media than from academic publications. That Wikipedia’s COVID-19 articles were based on respected sources in both the academic and popular media was found to be true even as the pandemic and number of articles about it grew. Our investigation further demonstrates that despite a surge in preprints about the virus and their promise of cutting-edge information, Wikipedia preferred published studies, giving a clear preference to open-access studies. A temporal and network analysis of COVID-19 articles indicated that remaining up-to-date did come at a cost in terms of quality, but also showed how preexisting content helped regulate the flow of new information into existing articles. In future work, we hope the tools and methods developed here in regards to the first wave of the pandemic will be used to examine how these same articles fared over the entire span of 2020, as well as helping others use them for research into other topics on Wikipedia. We observed how Wikipedia used volunteer-editors to enforce a rigid sourcing standards - and future work may continue to provide insight into how this unique method can be used to fight disinformation and to characterize the knowledge infrastructure in other arenas.

## Acknowledgments

J.S. is a recipient of the Placide Nicod foundation, and R.A. is a recipient of the Azrieli Foundation fellowship. We are grateful for their financial support.

## Supplementary information

**Table 1.**
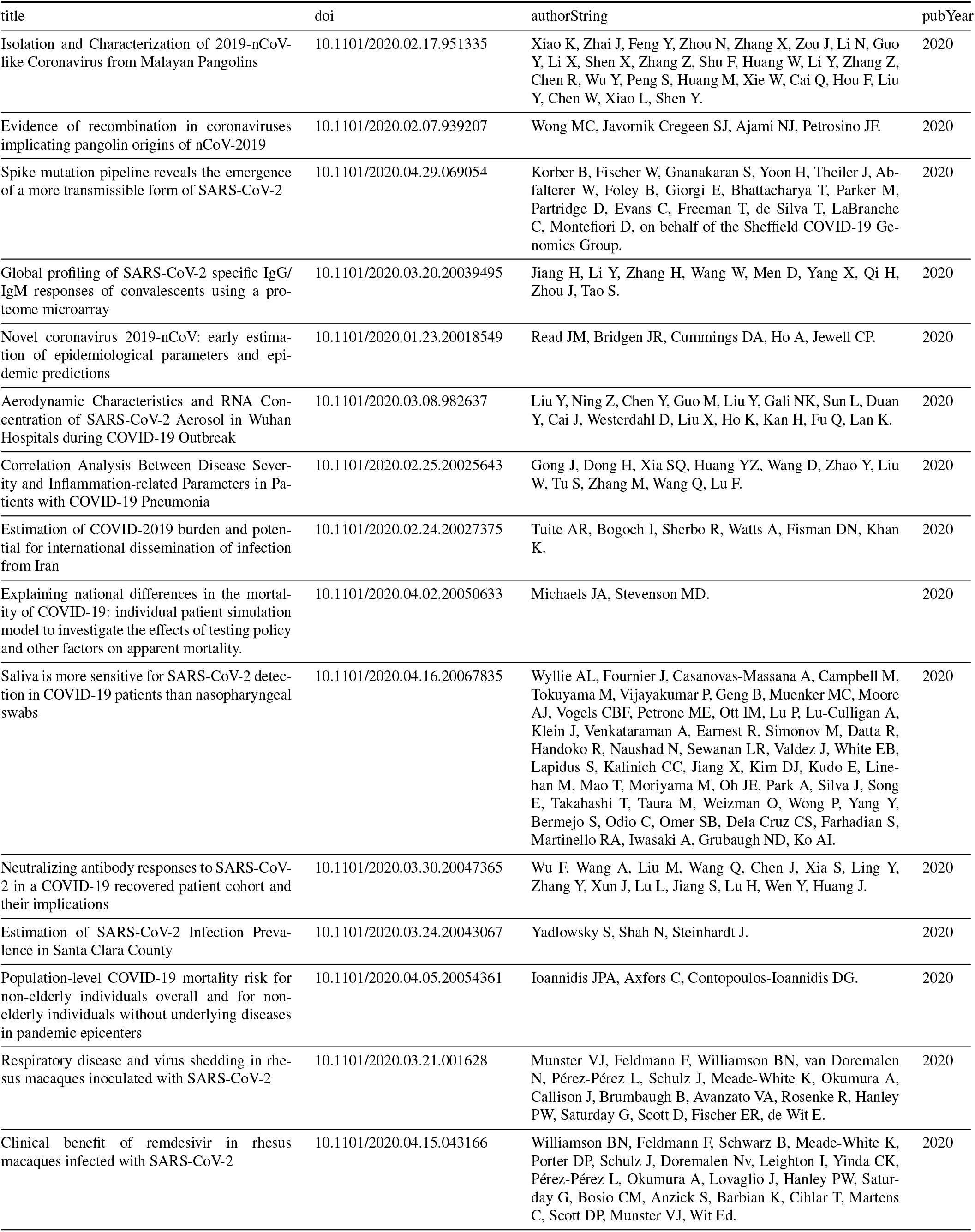
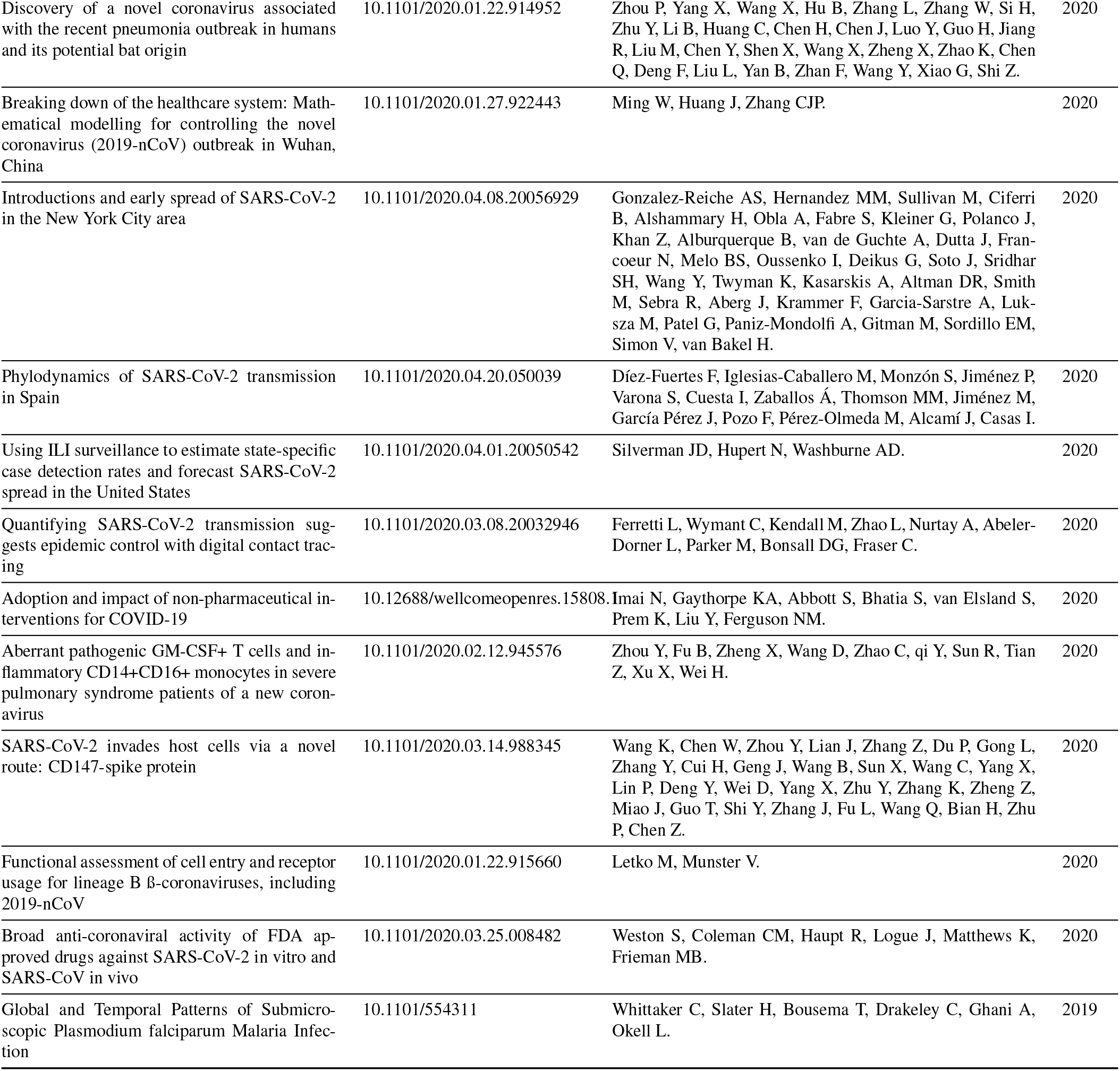
Preprints cited within the Wikipedia COVID-19 Corpus

**Table 2.**
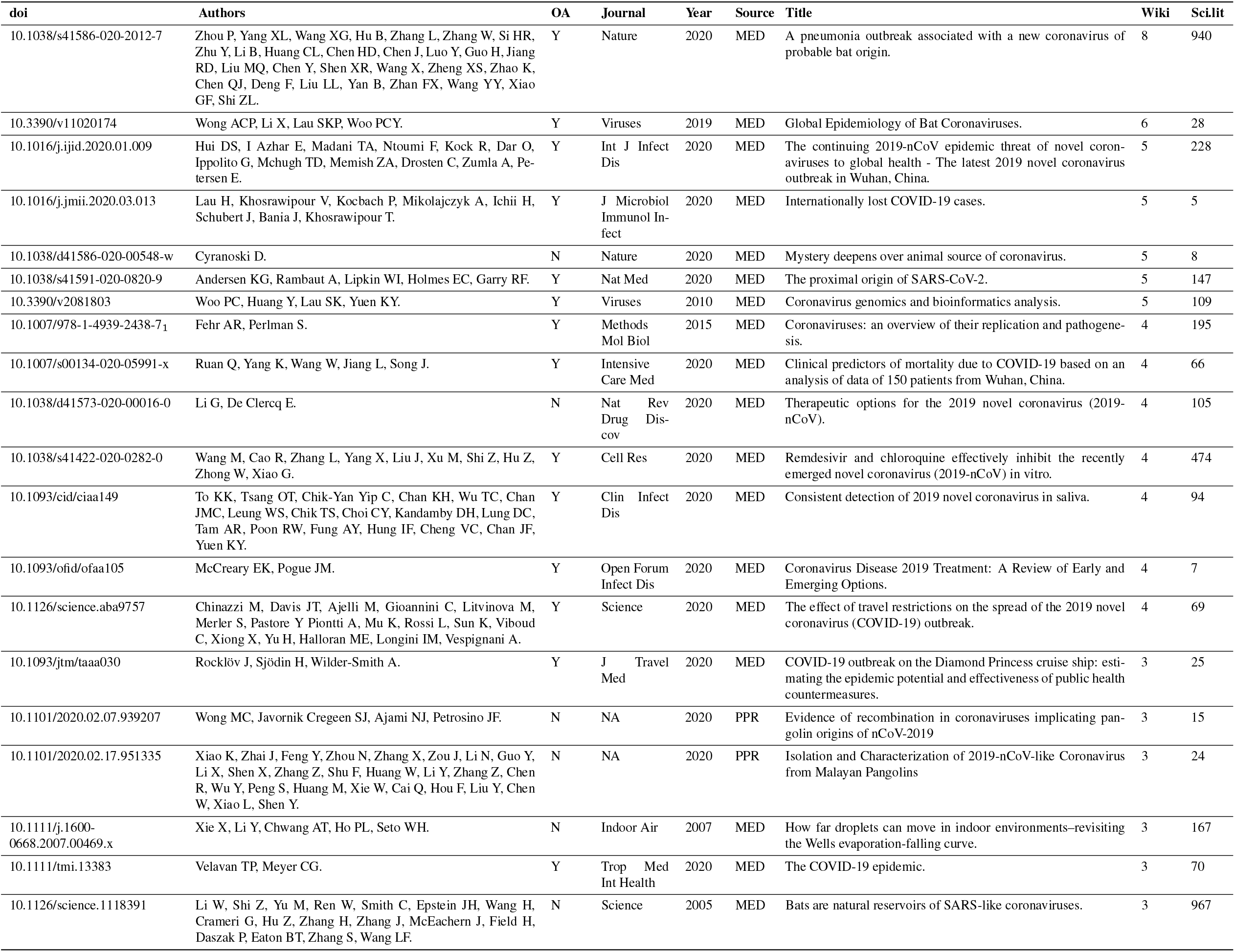
Most cited scientific papers in COVID-19 Wikipedia corpus

**Table 3.**
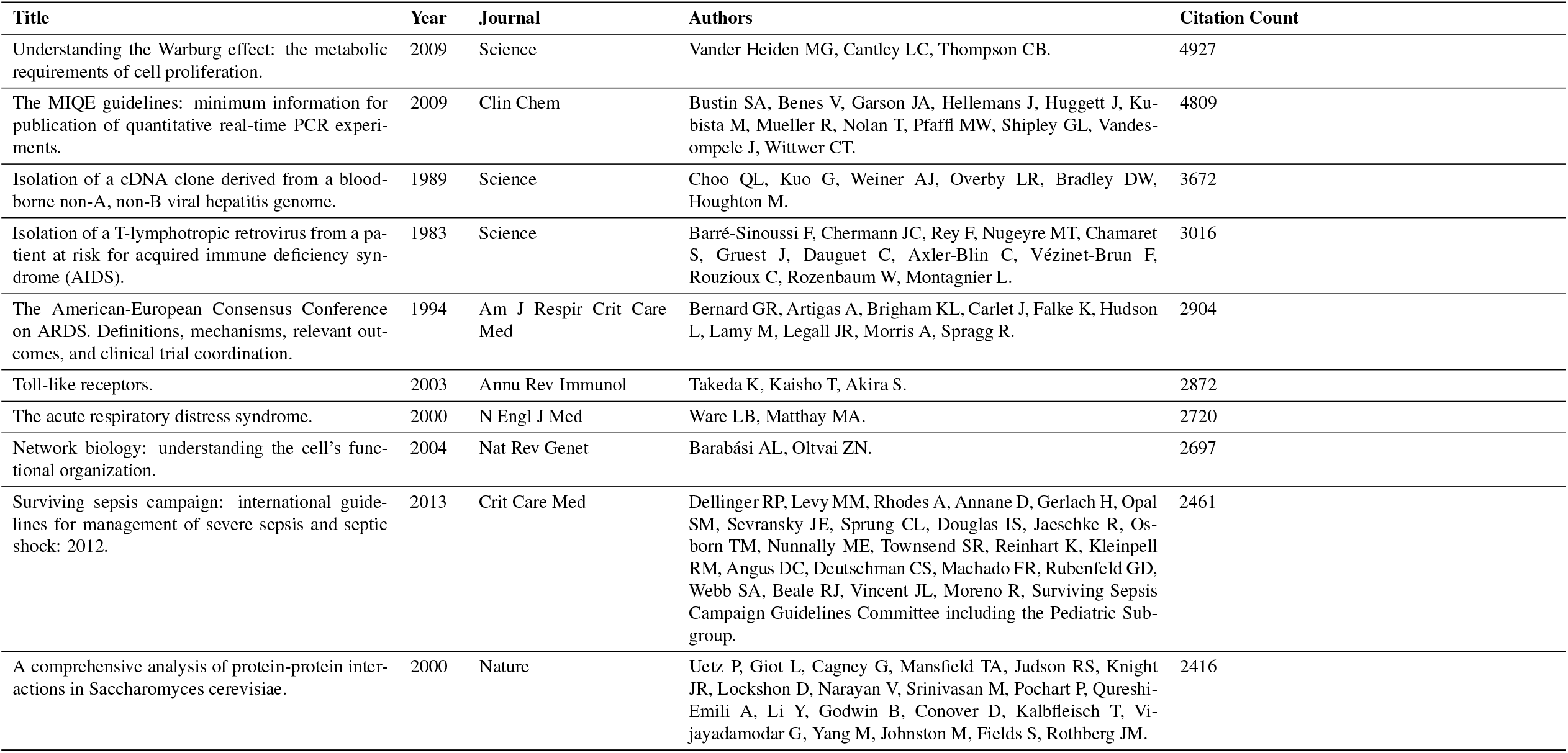
Most cited scientific papers in the scientific literature within COVID-19 Wikipedia corpus

## SI datasets

1. Table of scientific paper form europmc COVID-19 cited in wikipedia
2. Table of Wikipedia article-DOI network
3. Table of protected wikipedia COVID-19 articles

---

**Fig. S1.**
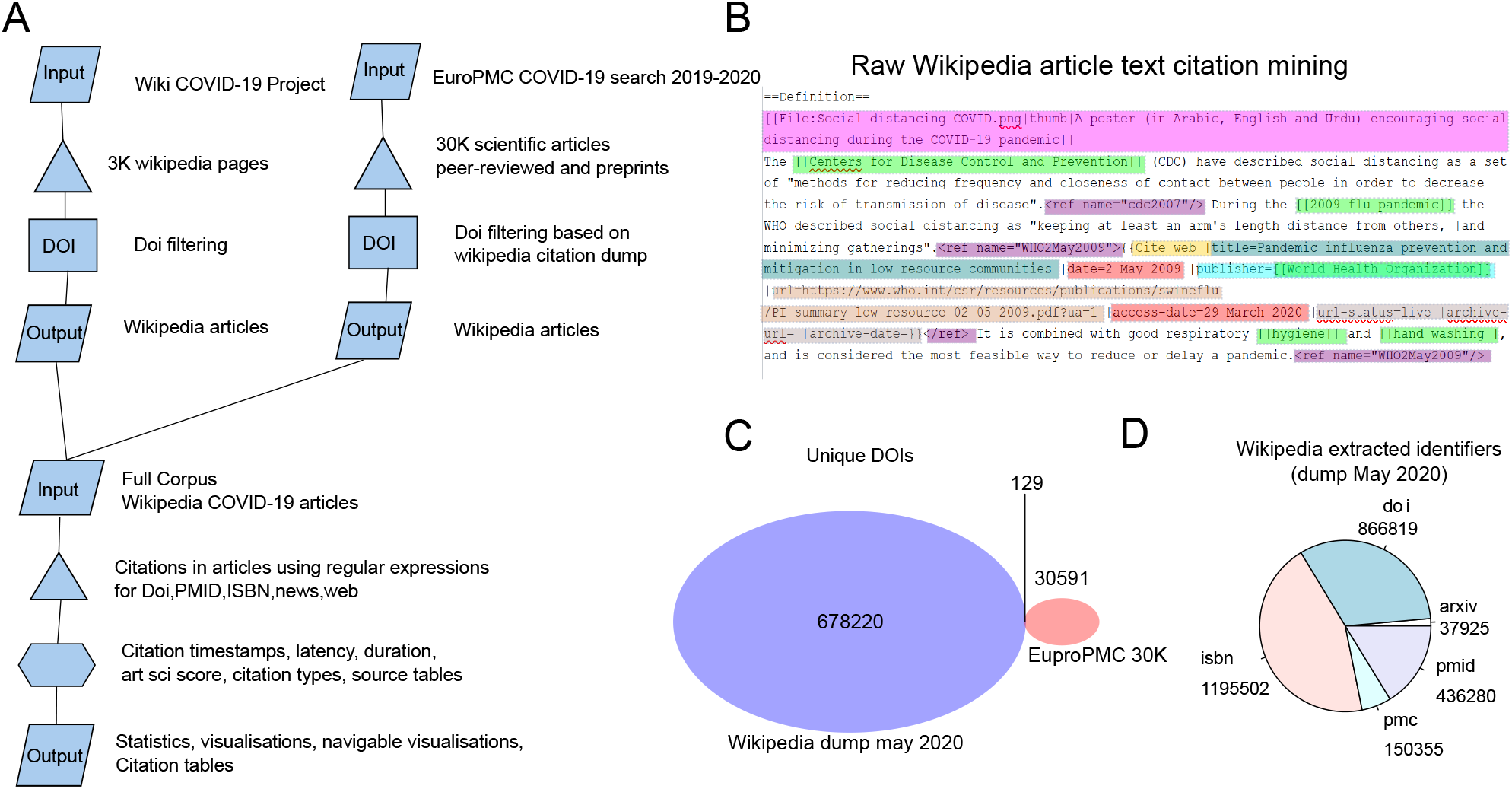
Corpus identification and citation extraction pipeline. A) Scheme of the Corpus delimitation rational and citation extraction. To delimit our corpus of Wikipedia articles containing Digital Object Identifier (DOI), we applied two different strategies. First we scraped every Wikipedia pages form the COVID-19 Wikipedia project (about 3K pages) and we filtered them to keep only page containing DOI citations (149 Wikipedia articles). For our second strategy, we made a search with EuroPMC on COVID-19, SARS-CoV2, SARS-nCoV19 (30,000 sci papers, reviews and preprints) and a selection on scientific papers form 2019 onwards that we compared to the Wikipedia extracted citations from the English Wikipedia dump of May 2020 (860’000 DOIs). This search led to 91 Wikipedia articles containing at least one citation of the EuroPMC search. Taken together, from our 231 Wikipedia articles corpus we extracted DOIs, PMIDs, ISBNs, websites and URLs using a set of regular expressions, as described in the methods. Subsequently, we computed several statistics for each Wikipedia article and we retrieved Atmetics, CrossRef and EuroPMC information for each DOI. Finally, our method allows to produce tables of citations annotated and extracted information in each Wikipadia articles such as books, websites, newspapers. In addition, a timeline of Wikipedia articles and a network of Wikipedia articles linked to scientific papers is built. B) Example of raw Wikipedia text from the social distancing article highlighted with several parsed items from a reference. pink: a hyperlink to an image file, green: Wikipedia hyperlinks, purple: reference, yellow: citation type, dark green: citation title, red: citation date, orange: citation URL. C) Overlap between DOI from the Wikipedia dump and the 30K EuroPMC COVID-19-related scientific articles and preprints D) number of extracted citations with *mwcite* from the English Wikipedia dump of May 2020.

**Fig. S2.**
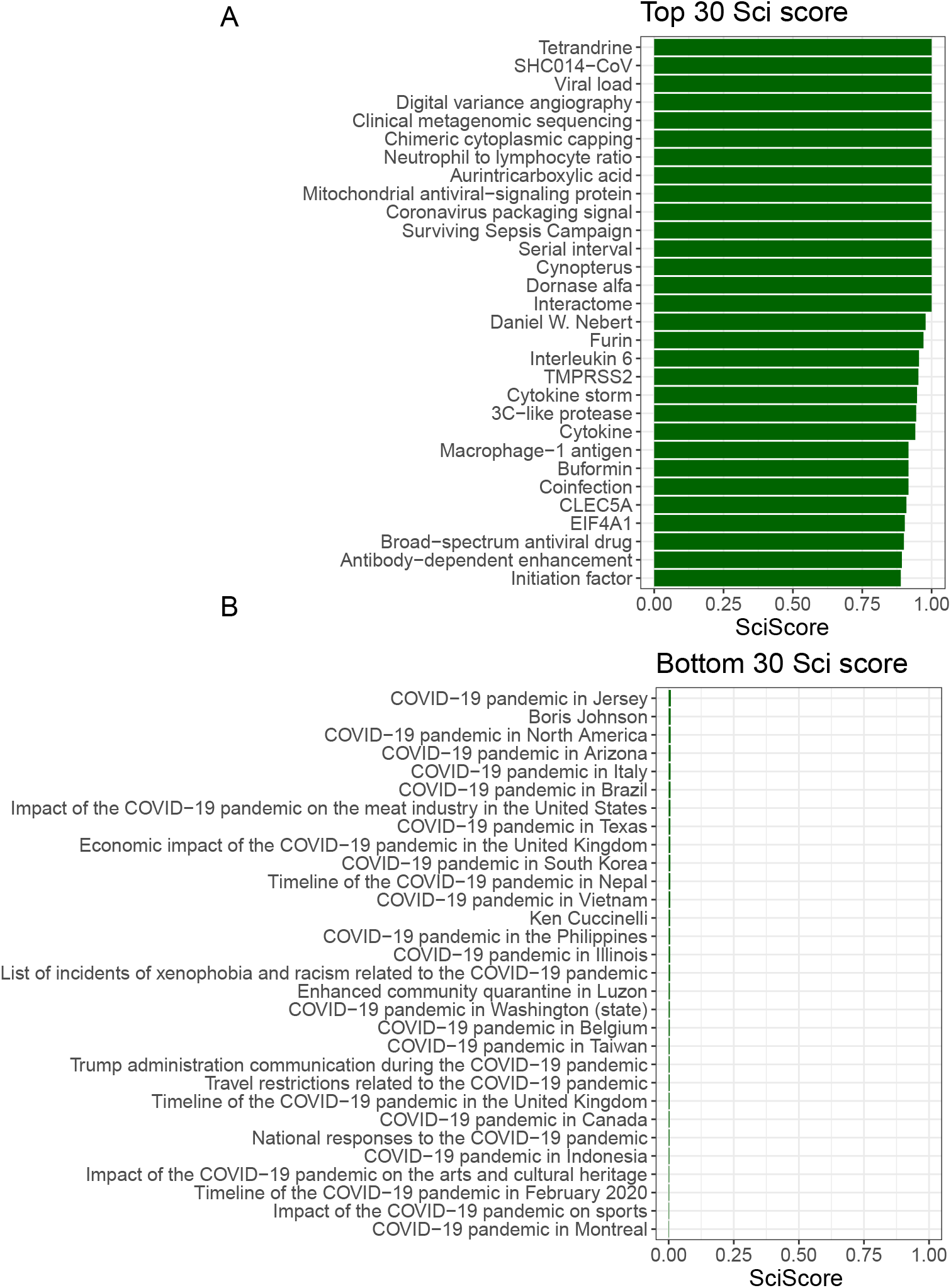
Top and bottom scientific score from Wikipedia article COVID-19 corpus. The scientific score was computed based on the reference content of each Wikipedia article from the COVID-19 corpus as defined in the methods section. A) Top 30 scientific article from the COVID-19 corpus. B) Bottom scientific article from the COVID-19 corpus.

**Fig. S3.**
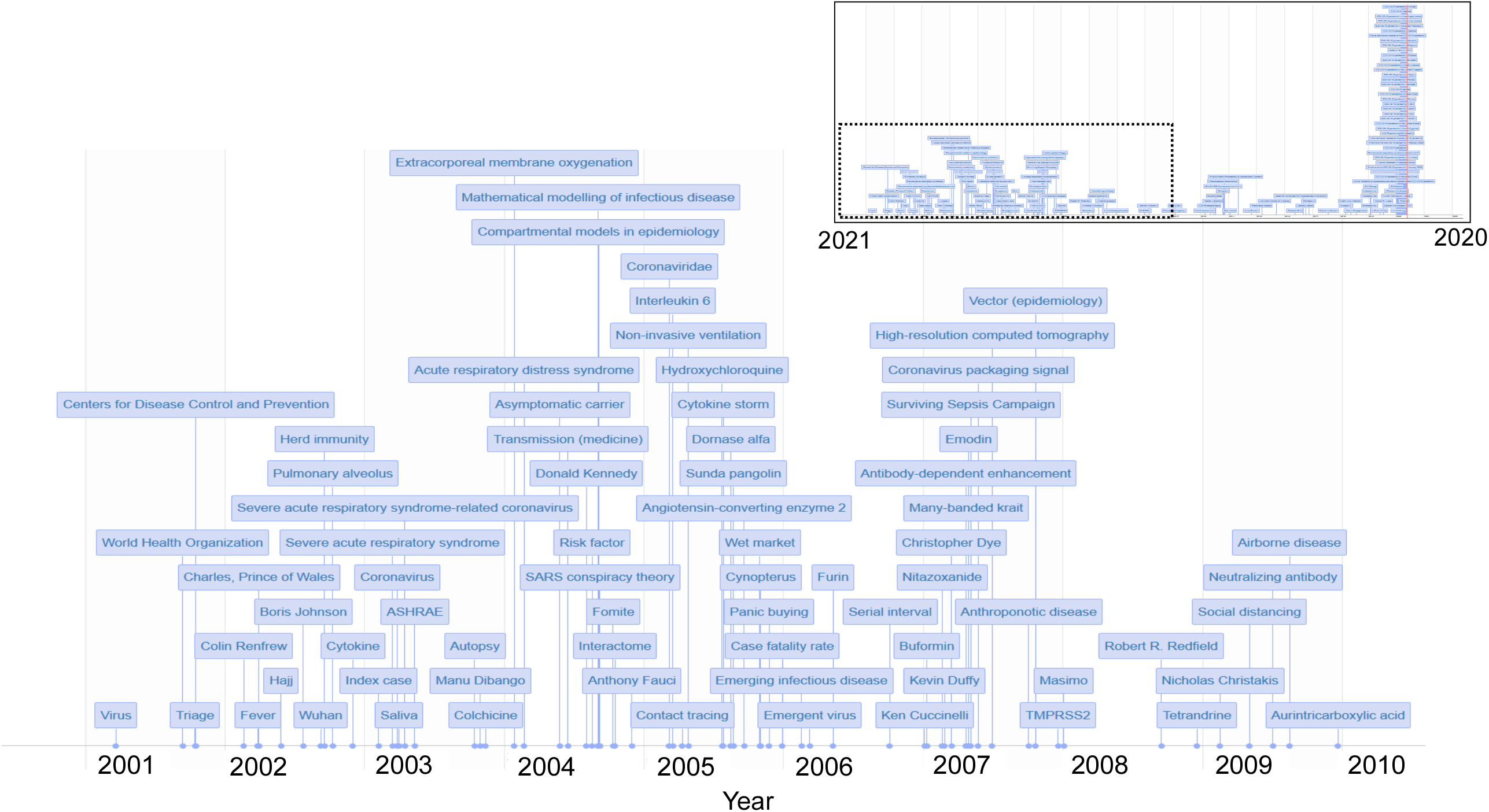
Timeline of the Wikipedia COVID-19 corpus. See here for an interactive version of the timeline.

**Fig. S4.**
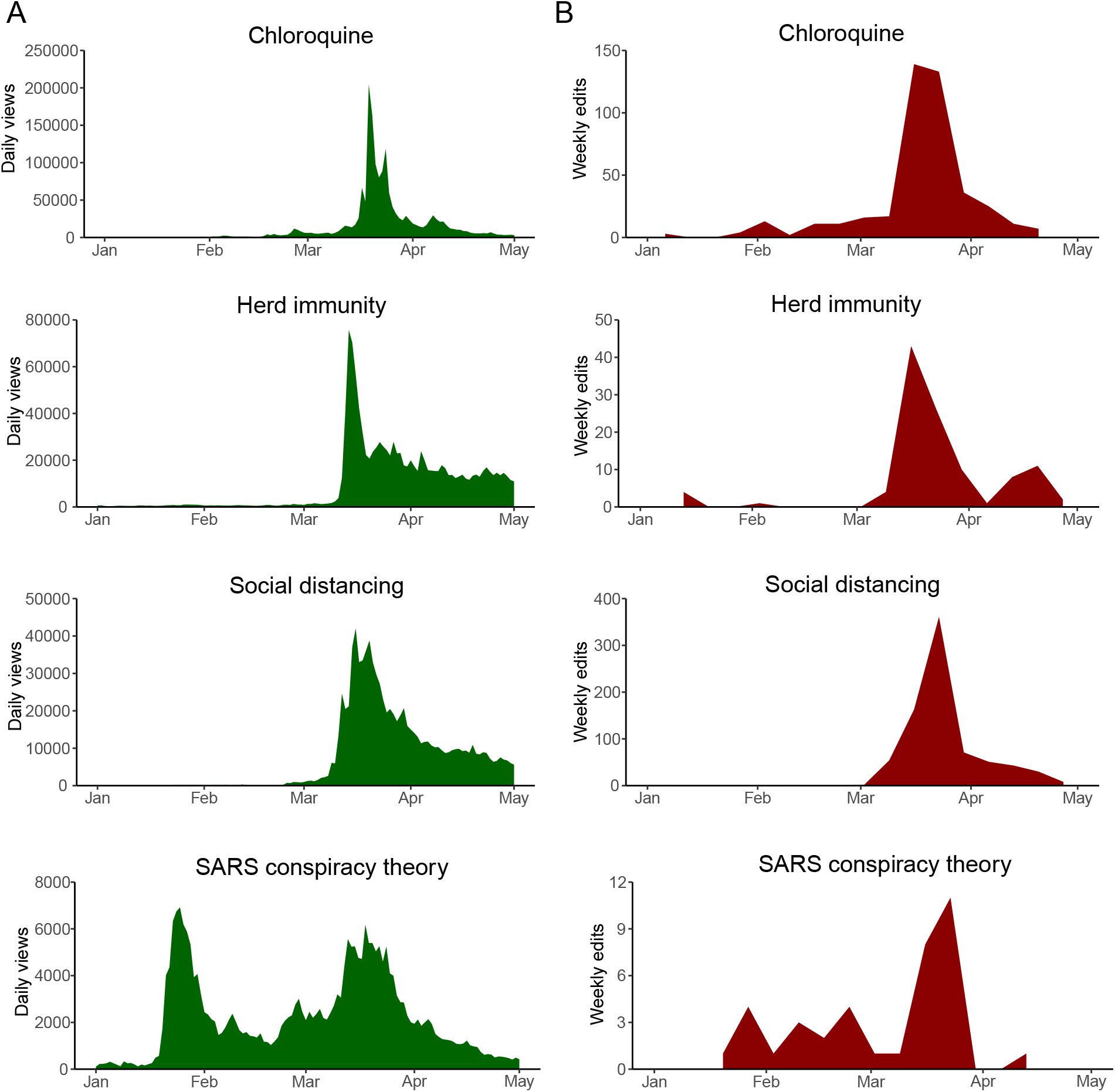
Wikipedia article page views and edits during COVID-19 pandemics. A) Daily page views and B) weekly edits for selected Wikipedia articles.

**Fig. S5.**
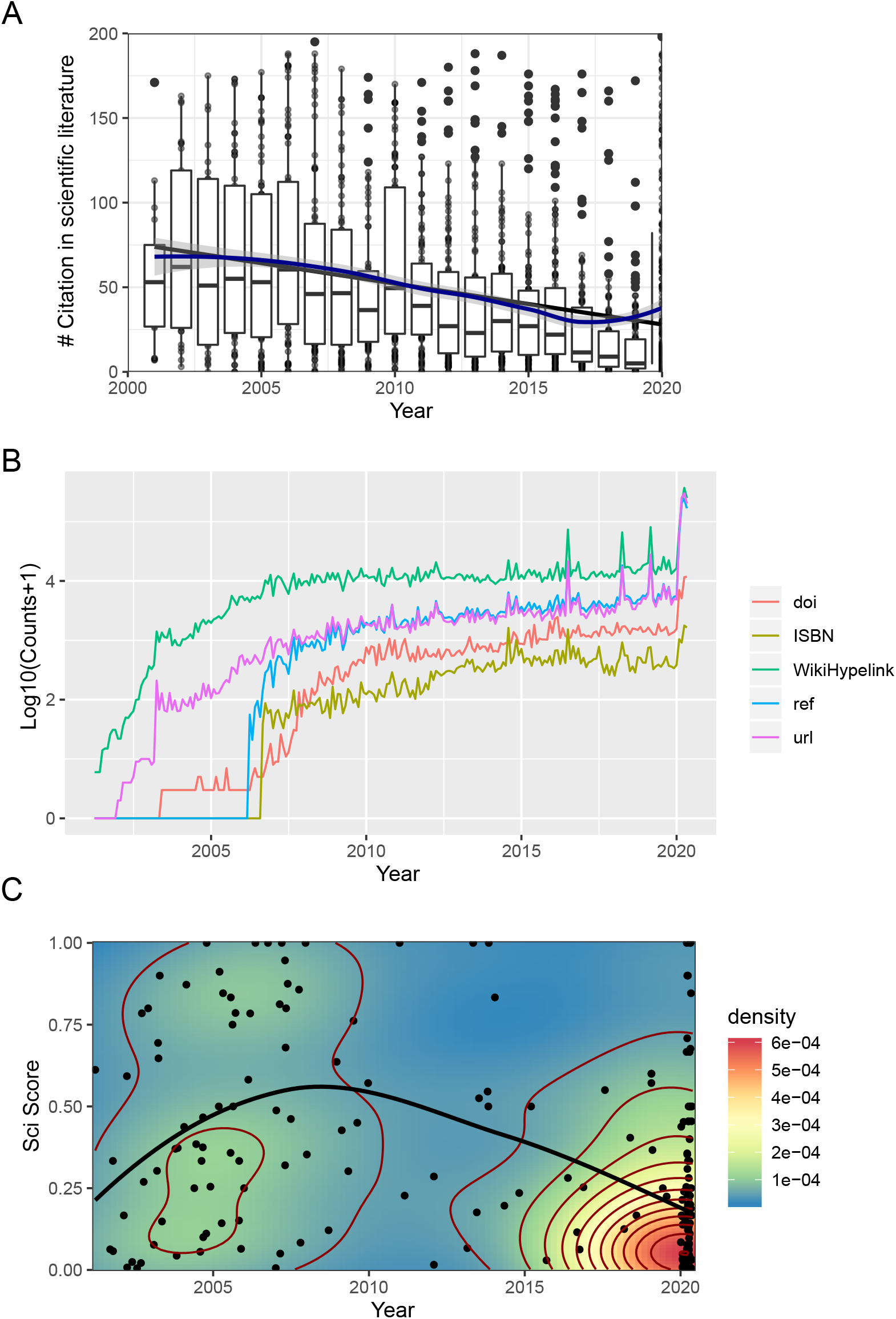
Historical perspective, citations count, citation type and scientific score of the COVID-19 corpus. A) Scientific literature citation count in function of the year of publication. B) Citation count in function of the year for different type of citation (doi, isbn, hyperlink, url). C) Scientific score in function of the creation date of wikipedia article.

## Notes

https://zenodo.org/record/3901741#.YD0Pdnk6-cw

https://github.com/jsobel1/WikiCitationHistoRy

### Competing Interest Statement

The authors have declared no competing interest.

### Summary of Updates

Revised version after first round of review from GigaScience.

